# Is the population of the Critically Endangered White-bellied Heron in Namdapha Tiger Reserve, India, declining?

**DOI:** 10.1101/2024.05.13.593898

**Authors:** Rohan K. Menzies, Kulbhushansingh R. Suryawanshi, Rohit Naniwadekar

## Abstract

The Critically Endangered White-bellied Heron *Ardea insignis* is a large-bodied, range-restricted, piscivorous bird found in the Himalayan mountains. With fewer than 60 individuals remaining globally, regular monitoring efforts are required to understand the conservation status of this rare species better. Here, we present the results of river surveys from 2017 and 2023-24 for the White-bellied Heron in an important protected area, the Namdapha Tiger Reserve, in Arunachal Pradesh. The encounter rate in 2017 was 0.55 herons per km, however; we did not detect the bird during intensive river sampling in the same stretches in 2023-24. This area within the reserve was once a stronghold for the species and they currently appear to be declining or locally extinct. Given the declines within this Protected Area, the drivers for the same need to be determined. While repeated monitoring of rivers has helped reveal this decline, locating and conserving stretches of river utilised by the remaining individuals away from the core zone is now critical.

[Supplementary material for this article can be found at https://doi.org/xxx]

---

The White-bellied Heron *Ardea insignis* is the rarest heron species in the world (Weseloh et al., 2022). This large, Critically Endangered river bird is estimated to have approximately 60 individuals worldwide (Stanley Price & Goodman, 2015). White-bellied Herons are now extinct in Bangladesh and Nepal, and their global population is declining. Their extant distribution is extremely patchy across Bhutan, Myanmar, and a few locations in northeast India (Stanley Price & Goodman, 2015). Existing studies on the heron have focussed on their foraging (Mondal & Maheswaran, 2021), breeding (Mondal & Maheswaran, 2014; Acharja, 2020, Reddy et al., 2021), or anecdotal natural history observations (Maheshwaran, 2008; Acharja et al., 2021; Khandu, 2022). Information on encounter rates or densities of heron that can aid population monitoring is relatively limited (but see Menzies et al., 2021). Since 2016, herons in Bhutan have stopped re-using nests each year and have changed to a single-use nesting pattern while abandoning older nesting areas, leading to unsuccessful nests in many locations (Acharja, 2020). There are now as few as five breeding pairs in Bhutan (Acharja, 2020) and two active nests recorded from two locations in Arunachal Pradesh (Mondal, 2018; Reddy et al., 2021). Given their rarity and critical conservation status, it is imperative to monitor their populations.

There are two known populations -the western population in Bhutan and the eastern population in the Indo-Myanmar region. The eastern part of Arunachal Pradesh state in India, at the junction of the Eastern Himalaya and the Indo-Myanmar region, is an important region for the eastern population of the White-bellied Heron. Given the political instability in Myanmar, the eastern Arunachal region is among the last refuge for the eastern population of the heron. In this region, the Namdapha Tiger Reserve (area: 1,985 km^2^), which has harboured low but stable heron populations in this region (Maheshwaran, 2008; Mondal, 2018; Menzies et al. 2021), is the largest and most important protected area for the species in the region. Recently, the bird has been reported from areas adjoining Namdapha (Habung et al., 2019; Reddy et al., 2021). A previous systematic survey of the heron in the central and western parts of Namdapha (Menzies et al., 2021) provided a unique opportunity to determine their status in Namdapha. Here, we demonstrate a significant decline in the White-bellied Heron encounter rates from the central and western parts of Namdapha based on river surveys in 2017 and 2023-24.

We sampled the Deban, Namdapha, and Noa-Dihing Rivers in Namdapha TR (Fig. 1). In 2017, we surveyed seven transects in November ranging from 600 m to 2 km with a total effort of 12.6 km, as part of the Eastern Himalayan-wide survey of the White-bellied Heron (Menzies et al., 2021). In February 2023, we sampled five transects ranging from 3.8 km to 7.7 km with a total effort of 29.7 km, and between November 2023 and March 2024, we surveyed 18 transects of 500 m length five times each with a total effort of 45 km. We recorded all sightings (perched or flying) of the White-bellied Heron during and outside the transect walks. Additional sampling details are in Table 1.

**TABLE 1.** A comparison of White-bellied Heron detections and encounter rates from line transect surveys in Namdapha Tiger Reserve, Arunachal Pradesh, India intermittently from 2014-2024.

| Survey | Duration | No. of Transects | Average Length | Total Effort | Total Detections | Total Individuals | Encounter Rate (per km) |
| --- | --- | --- | --- | --- | --- | --- | --- |
| Mondal 2018 | November 2014 - January 2017 | 65 | 0.62 km | 40.0 km | 7 | 8 | 0.20<sup>1</sup> |
| Menzies et al. 2021 | November 2017 | 7 | 1.80 km | 12.6 km | 6 | 7 | 0.55 |
| This study | February 2023 | 5 | 5.93 km | 29.7 km | 0 | 0 | 0 |
| This study | November 2023 - March 2024 | 90 | 0.50 km | 45.0 km | 0 | 0 | 0 |

<sup>1</sup>The author used detections/effort to calculate encounter rate which is 0.175 but we used total individuals/effort in 2021, so have converted this to 0.2.

**FIG. 1.**
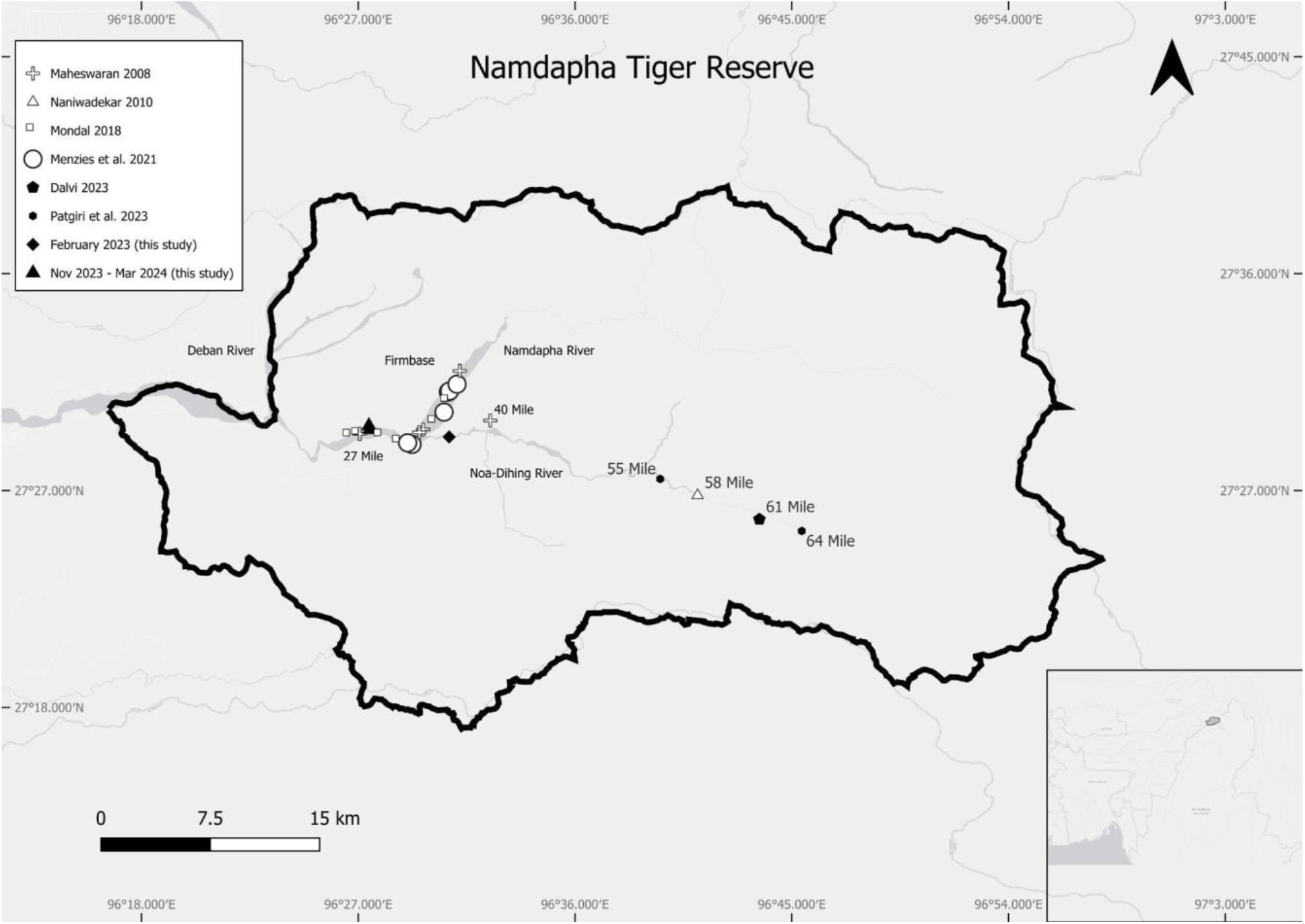
Sighting locations of the White-bellied Heron Ardea insignis in Namdapha Tiger Reserve from 2005 to 2024. The white symbols denote sightings before 2020 and black symbols denote records from 2021-2024.

In 2017, we detected seven herons during sampling and the mean (SE) encounter rate of the White-bellied Heron was 0.55 (± 0.24) individuals per km. However, in 2023 and 2024, we did not detect a single heron during sampling but opportunistically recorded one individual in February 2023 and two sightings between November 2023 and March 2024. In February 2023, one individual was spotted between 38 Mile and Firmbase. Between November 2023 to March 2024, a single individual was sighted twice near Bor Nala.

The earliest surveys of the White-bellied Heron along the entire stretch of Noa-Dihing River till Vijaynagar reported all five sightings of the heron from below 40 Mile (Maheshwaran, 2008). There have been incidental reports of the heron from sites above 40 Mile till 65 Mile from 2007 onwards. The heron was spotted in the 52 Mile area by Umesh Srinivasan and Shashank Dalvi (pers. comm.) in 2007 (Srinivasan et al., 2010). RN photographed the bird at 58 Mile in February 2010 (https://media.ebird.org/catalog?userId=USER493606&taxonCode=whbher2&mediaType=photo). Subsequently, Patgiri et al. (2023), have reported the heron from the 64 Mile area further east (Fig. 1). Earlier, Firmbase was the preferred site for heron sightings by bird enthusiasts (pers. obs.). However, in the recent past, due to better accessibility provided by the newly constructed Miao-Vijaynagar Road and reduced sightings of the heron at Firmbase area, bird enthusiasts prefer to spot it around the 65 Mile area (pers. obs. and Shashank Dalvi, pers. comm.). Some locals suggest the movement of the bird from Firmbase to the higher reaches of Noa-Dihing in the eastern part of the park. Unfortunately, there is only incidental observation data on the heron from the eastern part of Namdapha. This precludes us from ascertaining whether the birds have moved from the lower reaches to the higher reaches or whether there has been a reduction in the population of herons in central and western Namdapha.

Our temporally replicated surveys for the White-bellied Heron at Namdapha TR, a site critical for heron conservation, demonstrate declines in the heron population in the central and western parts of Namdapha. Patgiri et al. (2023) also failed to detect the heron in the Firmbase area; however, they reported sighting it at the confluence of Namdapha and Noa-Dihing. Our sampling effort for river birds in Namdapha, especially between November 2023 and March 2024, is comparable to other such efforts for river birds in the Himalaya (Sinha et al., 2019). We are uncertain about the reasons behind these declines in the western and central parts of Namdapha. Previous studies have suggested anthropogenic disturbances as a possible driver for the reduction in encounters of the heron in the central and western parts of Namdapha (Patgiri et al., 2023). Based on the anthropogenic activities recorded during the 2017 and 2023-24 surveys, there is a slight reduction in human use in the protected area. When comparing the number of 100 m segments within a site with signs of fishing, we found evidence of fishing in 5.6% of segments in 2017 (Menzies et al., 2021) versus just 1.8% of segments in 2023-24 despite having nearly four times the effort. Similarly with human presence, in 2017, it was observed in 9.5% of segments (Menzies et al., 2021) and 7.6% of segments in 2023-24, respectively. The sampled riverine habitat inside the Protected Area has remained mostly unaltered, and the local human activity (except activity by villagers staying around Firmbase) along the river has likely reduced, given the recent road connectivity between Miao and Vijaynagar. In February 2023, we detected a school of dead fish likely killed by dynamiting at Firmbase. However, during our surveys from November 2023 to March 2024, we did not detect any evidence of destructive fishing methods inside Namdapha. Our understanding of temporal changes in fish abundance, a potentially important driver of White-bellied Heron abundance, is poor. Like in the past, fishing continues inside the park. Fish abundance needs to be compared between the eastern and western parts of Namdapha to determine if there are differences in abundance and body size of the heron’s preferred prey fish.

In any case, there has been a reduction in the use of central and western parts of Namdapha either due to local extinctions or through the movement of birds away from the area. There is a need for systematic sampling of the eastern part of Namdapha so trends in the heron population in the eastern part of Namdapha can be determined. Moreover, systematic efforts to find and monitor heron’s active nests away from the core area is critical to determine their breeding success and the potential factors influencing that. As birdwatchers continue to frequent parts of eastern Arunachal Pradesh, new site records for the bird are reported. As part of the river bird study, we sampled areas outside Namdapha Tiger Reserve and five transects in Kamlang Tiger Reserve (Menzies et al., unpublished data); however, we did not detect the heron. Systematic surveys and long-term monitoring in more remote parts of Kamlang and the higher reaches of Anjaw District are critical to understanding whether the birds are using these sites for nesting. Despite the apparent reduction in White-bellied Herons at the core of Namdapha Tiger Reserve, it is important to regularly monitor the remaining individuals as it could be the last hope for this species on the verge of extinction. A clear understanding of the reasons for the decline, which is yet unknown, would benefit the conservation management of the critical eastern population.

## Author contributions

Study design: RKM, RN, KRS; fieldwork: RKM, RN; writing: RKM, RN, KRS.

## Acknowledgements

We are grateful to Japang Pansa, Dhan Bahadur Limbu, Laiphung Wangnow and Phupla Singpho for assisting with the fieldwork. We thank Vidyadhar Atkore for useful discussions. We thank the Rufford Small Grants for Nature Conservation (37739-1) and Rohini Nilekani Philanthropies for funding this project. We thank the Arunachal Pradesh Forest Department for providing the necessary research permits (CWL/GEN/2018-2019/Pt.IX/NG/352-57) to conduct this work.

## Ethical standards

This research abided by the *Oryx* guidelines on ethical standards. We received ethical clearance from the Research Ethics Committee of the Nature Conservation Foundation prior to the fieldwork.

---

**PLATE 1.**
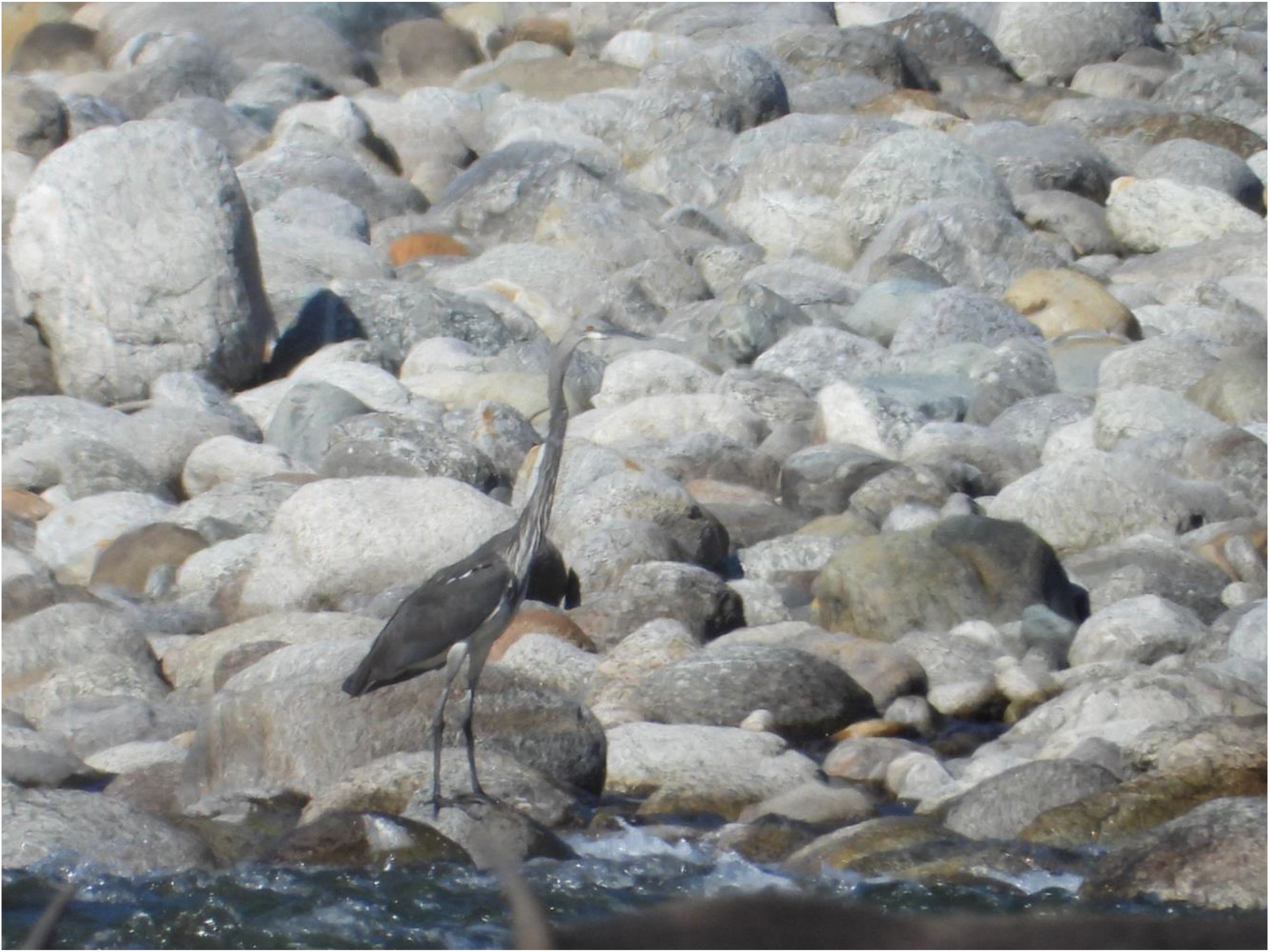
A White-bellied Heron *Ardea insignis* photographed in November 2023 along the Noa-Dihing River in Namdapha Tiger Reserve, Arunachal Pradesh, India. This individual was spotted outside of our fixed transects that were monitored five times between November 2023 to March 2024.

